# Development of the Functional Connectome Topology in Adolescence: evidence from Topological Data Analysis

**DOI:** 10.1101/2021.10.04.463103

**Authors:** Zeus Gracia-Tabuenca, Juan Carlos Díaz-Patiño, Isaac Arelio, Martha Beatriz Moreno, Fernando A. Barrios, Sarael Alcauter

**Affiliations:** Universidad Nacional Autónoma de México, Instituto de Neurobiología, Querétaro, Mexico; Universidad Nacional Autónoma de México, Instituto de Matemáticas, Querétaro, Mexico

## Abstract

Adolescence is a crucial developmental period in terms of behavior and mental health. Therefore, understanding how the brain develops during this stage is a fundamental challenge for neuroscience. Recent studies have modelled the brain as a network or connectome, mainly applying measures from graph theory, showing a change in its functional organization such as an increase in its segregation and integration. Topological Data Analysis (TDA) complements such modelling by extracting high-dimensional features across the whole range of connectivity values, instead of exploring a fixed set of connections. This study enquiries into the developmental trajectories of such properties using a longitudinal sample of typically developing participants (N = 98; 53/45 F/M; 6.7-18.1 years), applying TDA into their functional connectomes. In addition, we explore the effect of puberty on the individual developmental trajectories. Results showed that compared to random networks, the adolescent brain is more segregated at the global level, but more densely connected at the local level. Furthermore, developmental effects showed nonlinear trajectories for the integration of the whole brain and fronto-parietal networks, with an inflection point and increasing trajectories after puberty onset. These results add to the insights in the development of the functional organization of the adolescent.

**Significance Statement:** Topological Data Analysis may be used to explore the topology of the brain along the whole range of connectivity values instead of selecting only a fixed set of connectivity thresholds. Here, we explored some properties of the topology of the brain functional connectome, and how they develop in adolescence. First, we show that developmental trajectories are nonlinear and better explained by the puberty status than chronological age, with an inflection point around the puberty onset. The greatest effect is the increase in functional integration for the whole brain, and particularly for the Fronto-Parietal Network when exploring functional subnetworks.

## 1. Introduction

Adolescence is a critical development period with substantial impact on body and behavior. Particularly, the brain undergoes structural and functional changes that are influenced by pubertal hormones (Vijayakumar et al., 2018; Laube et al., 2020). Moreover, these changes occur along with a consolidation of cognitive and executive performance (Baum et al., 2017; Chai et al., 2017).

These insights had been addressed modelling the brain as a complex network of interacting components, either at task or rest conditions (Biswal et al., 1995; Smith et al., 2009). In this framework, the functional connectome is described by its system properties in biologically plausible terms, mainly using measures from graph theory (Rubinov and Sporns, 2010). Nevertheless, other methods had recently been applied to address high-dimensional data such as Topological Data Analysis (TDA) (Sizemore et al., 2018; Expert et al., 2019). TDA models the connectome as a topological space and characterizes its interaction patterns as geometric features, allowing it to simplify complex structures at different scales (Giusti et al., 2016; Santos et al., 2019; Centeno et al., 2021). In particular, TDA applied to functional connectomes is not affected by the potential biases of connectivity thresholding nor brain segmentation (Lee et al., 2012; Gracia-Tabuenca et al., 2020).

In terms of functional organization of the brain, previous cross-sectional studies have shown that the adolescent period is characterized by an increase of the modularity and specialization (Fair et al., 2009; Satterthwaite et al., 2013a; Gu et al., 2015), with prominent effects in frontal and parietal systems, along with executive performance (Marek et al., 2015; Gracia-Tabuenca et al., 2021). However, as far as we are concerned, TDA in human connectomes have mainly been applied into neuropsychiatric disorders (Lee et al., 2012, 2017; Gracia-Tabuenca et al., 2020; Li et al., 2020), but not to characterize the typical development. There is still a huge degree of incertitude in this field due to the great variability between samples, sexes, and cultures (Sawyer et al., 2018), with special emphasis in the fact that some individuals have faster or slower pubertal development even when they have the same chronological age (Blakemore et al., 2010; Vijayakumar et al., 2018). To this regard, longitudinal trajectories and pubertal markers are highly valuable to describe adolescent development.

Therefore, this study focuses on characterizing the development of the functional connectome in the adolescent period applying TDA into a longitudinal sample of typically developing subjects. In addition, the effect of pubertal status and linear vs. nonlinear trends are tested as well.

## 2. Methods

### 2.1 Sample

A general invitation was sent to local schools describing the study protocols and the inclusion/exclusion criteria. Inclusion criteria consisted of a full-term gestation (more or equal than 37 weeks). Exclusion criteria included academic year repetition and any neurological or psychiatric disorder identified with the MINI semi-structured interview. Signed informed consent for parents and verbal assent for minors was required. The study protocols followed the ethical principles of the Declaration of Helsinki and were approved by the Institutional Ethics Board.

The sample consisted of 98 typically developing participants (53 females, 45 males; age range: 6.7 -18.1 years old). From those, 41 returned for a second session, and 16 for a third. Follow-ups occurred after 5 years and the second after 2 years, respectively.

### 2.2 Pubertal status assessment

Participants fulfilled the Pubertal Development Scale (PDS; Petersen et al., 1988). PDS averages the response of five self-reported questions about growth spurt in height, pubic hair, and skin change for both sexes; plus breast growth and menarche for females and facial hair growth and voice change for males. Responses are absence (1), first signs (2), evident (3), and finished (4) pubertal spurt. Those participants under 10 years old were set to PDS level 1, following similar values in previous studies (Hibberd et al., 2015; van Duijvenvoorde et al., 2019). In addition, 8 missing values (4 females) were estimated via Generalized Additive Mixed Model (GAMM) with age-sex interaction locally estimated scatterplot smoothing (LOESS) curves.

### 2.3 Imaging

For each session, participants underwent an MRI protocol including a whole-brain fMRI sequence plus high-resolution T1-weighted images for anatomical reference. After five ‘dummy’ volumes for scan stabilization, a total of 150 fMRI volumes were obtained using a gradient recalled T2* echo-planar imaging sequence (TR/TE = 2000/40 ms, voxel size 4×4×4 mm^3^). Participants were instructed to lay down, close their eyes, and not to fall asleep. In order to ease participants to remain awake, the fMRI scan was applied at the beginning of the MRI session and always in the morning. T1 images were obtained using a 3D spoiled gradient recalled (SPGR) acquisition (TR/TE = 8.1/3.2 ms, flip angle = 12.0, voxel size 1×1×1 mm^3^). All brain imaging was acquired with a 3T MR GE750 Discovery scanner (General Electric, Waukesha, WI), using an 8-channel-array head coil. However, 20 sessions were acquired with a 32-channel coil, thus a covariate was included in the subsequent analyses.

### 2.4 Preprocessing

Structural T1 volumes were denoised with non-local means (Manjón et al., 2010) and N4 bias field correction (Tustison et al., 2010). fMRI datasets were preprocessed using FSL v.5.0.6 (Jenkinson et al., 2012; RRID:SCR_002823). Preprocessing steps included slice timing, head motion correction, brain extraction, intensity normalization, confound regression, spatial normalization, and 0.01-0.08 Hz band-pass filtering.

Considering that the pediatric population tends to move more inside the scanner (Satterthwaite et al., 2012), we implemented a strident strategy of confounding variables regression (Satterthwaite et al., 2013b). 36 parameters were regressed out from the fMRI time series, including the six head-motion estimated parameters plus the average time series of the global signal, white matter, and cerebrospinal fluid. The derivatives of these nine variables were also added, and the quadratic terms of those eighteen. Additionally, the volumes with a framewise displacement (FD-RMS; Jenkinson et al., 2002) greater than 0.25 mm (“spikes”) were included as confounds as well. This approach overpowers other widely used motion-mitigation methods (Ciric et al., 2017; Parkes et al., 2018; Graff et al., 2020). Eighteen sessions with less than four minutes without spike-volumes were discarded (Satterthwaite et al., 2013b; Parkes et al., 2018), therefore, the final sample consisted of 89 participants (39 male, age range: 6.7–18.1 y.o.), of whom 37 and 11 had two and three longitudinal sessions, respectively.

In addition, fMRI datasets were co-registered to their T1 volume with six degrees of freedom, and then warped twice using nonlinear SyN transformation (Avants et al., 2008; RRID:SCR_004757) to a pediatric template (NIHPD4.5-18.5; Fonov et al., 2011) and then to the MNI-152 standard template.

### 2.5 Functional Connectomes

Brain networks were defined based on 264 regions of interest (ROIs) as nodes (Power et al., 2011). Pairwise edges were calculated through Pearson’s correlation between the average fMRI preprocessed signal of every pair of ROIs.

These ROIs consist of 5mm-radius spheres with high consistency in task and rest tested in large fMRI databases (Power et al., 2011). Moreover, this set of ROIs can be grouped in thirteen functional networks. This segmentation has been applied in numerous pediatric studies (Satterthwaite et al., 2013a; Gu et al., 2015; Marek et al., 2015; Chai et al., 2017; Ciric et al., 2017; Gracia-Tabuenca 2020, 2021).

### 2.6 Topological Data Analysis

The functional connectome can be modeled as a topological space by the means of the Rips complex, defined as Rips(F,□). *F* stands for the set of nodes (same as the connectome nodes) and □ stands for the filtration value, which is a positive number that indicates which nodes of *F* with lower distance than it are connected. Then, the set of connected nodes of the Rips complex varies as a function of □. Additionally, algebraic properties can be extracted from the Rips complexes, the so-called Betti numbers. Specifically, Betti numbers of order zero or Betti-0 (B_0_) accounts for the number of components (i.e., the sum of groups of connected and isolated nodes), Betti-1 (B_1_) accounts for the number of “holes” in the two-dimensional space between connected nodes (Figure 1), and so on (for extensive review on TDA, we suggest Edelsbrunner et al., 2000; Sizemore et al., 2019).

**Figure 1.**
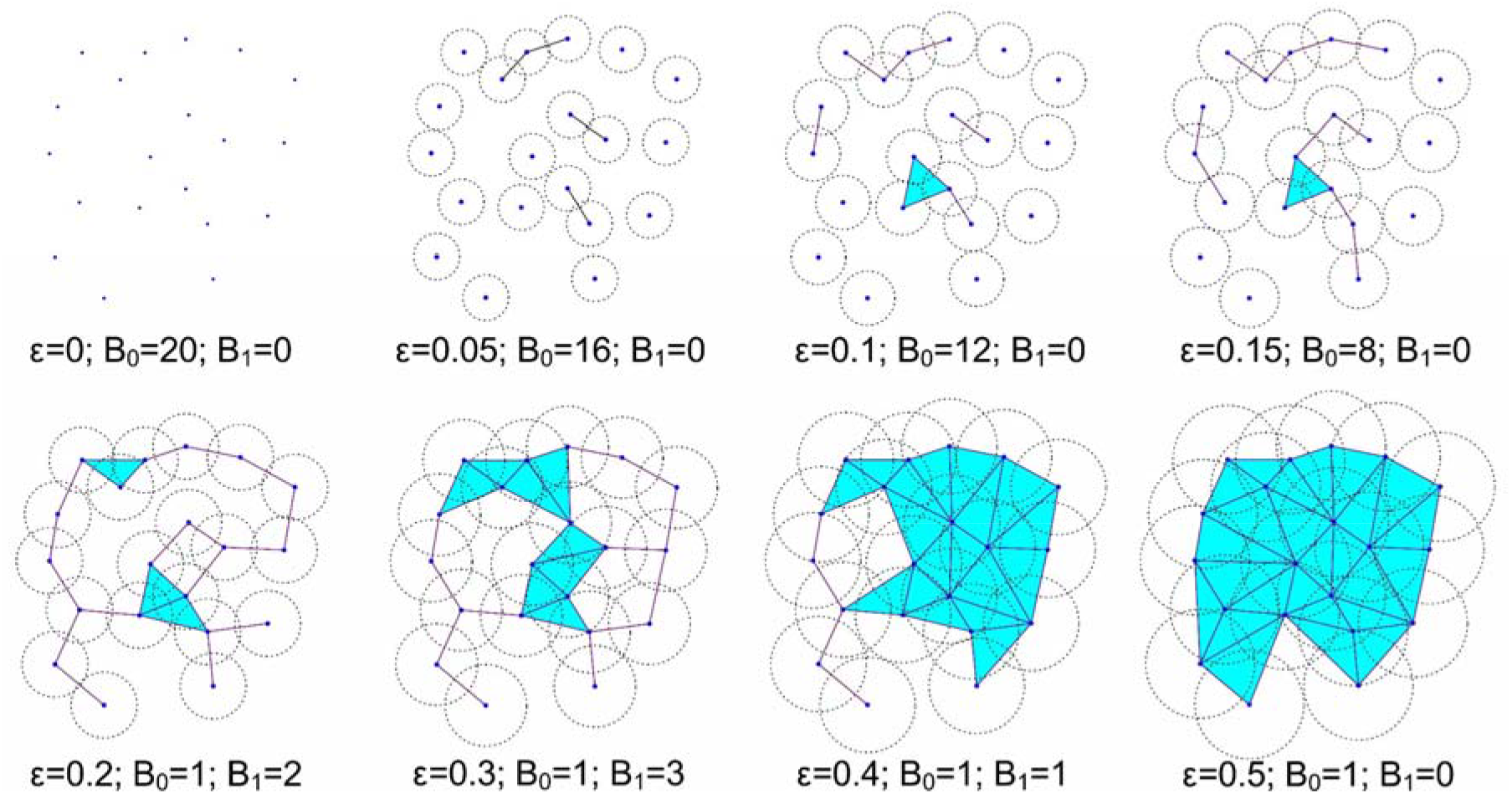
Betti-0 and Betti-1. Set of 20 nodes, eight filtration values ε, represented as the circle diameter and their corresponding Betti-0 (B_0_) and Betti-1 (B_1_). At ε = 0 the number of components (Betti-0) is equal to the number of nodes, even without links there are no holes. As the filtration value increases, the number of components (Betti-0) reduces while the number of holes (Betti-1) increases. A hole is a polygon with four or more sides, by definition a triangle is not a hole and if this is the case, we put a triangular surface (light blue). Eventually it will reach a single component containing all nodes and the number of holes becomes zero. Blue triangles are not holes as they are surfaces between triads of nodes.

In this study we focused exclusively on B_0_ and B_1_. As □ increases, the number of isolated nodes decreases in favour of the connected nodes, that is the B_0_ decreases and eventually will reach a single component where every node of *F* is connected (Figure 1). In contrast, at low values of □ there are no holes in the connected pattern of the topological space because there are not enough connections to build them. Similarly, at high values of □ the holes are “filled” because all pairwise connections within the component are accomplished. These holes represent serially distributed connections of nodes without shortcuts between them, while a filled hole means that those nodes are densely connected between them (Sizemore et al., 2018). Therefore, the greater amount of holes (i.e., B_1_) is reached at intermediate values of □. Both processes can be characterized by Betti curves as a function of □ (Figures 2 and 3).

**Figure 2.**
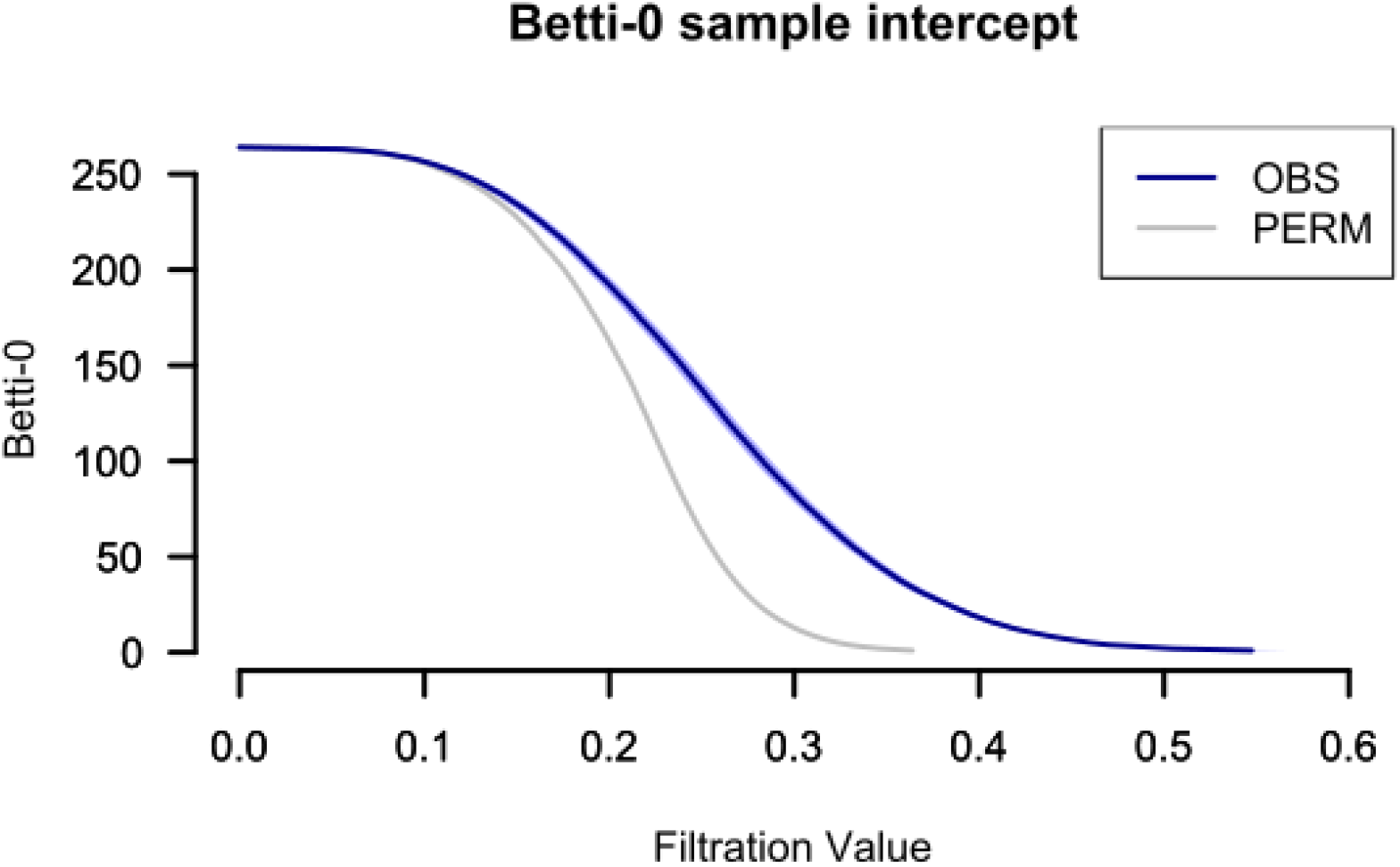
Sample intercept B_0_ curve (with 95% confidence interval) in blue. Average of 1000 bootstrapped connectomes with random edge-rewiring B_0_ curve (with 95% confidence interval) in gray.

**Figure 3.**
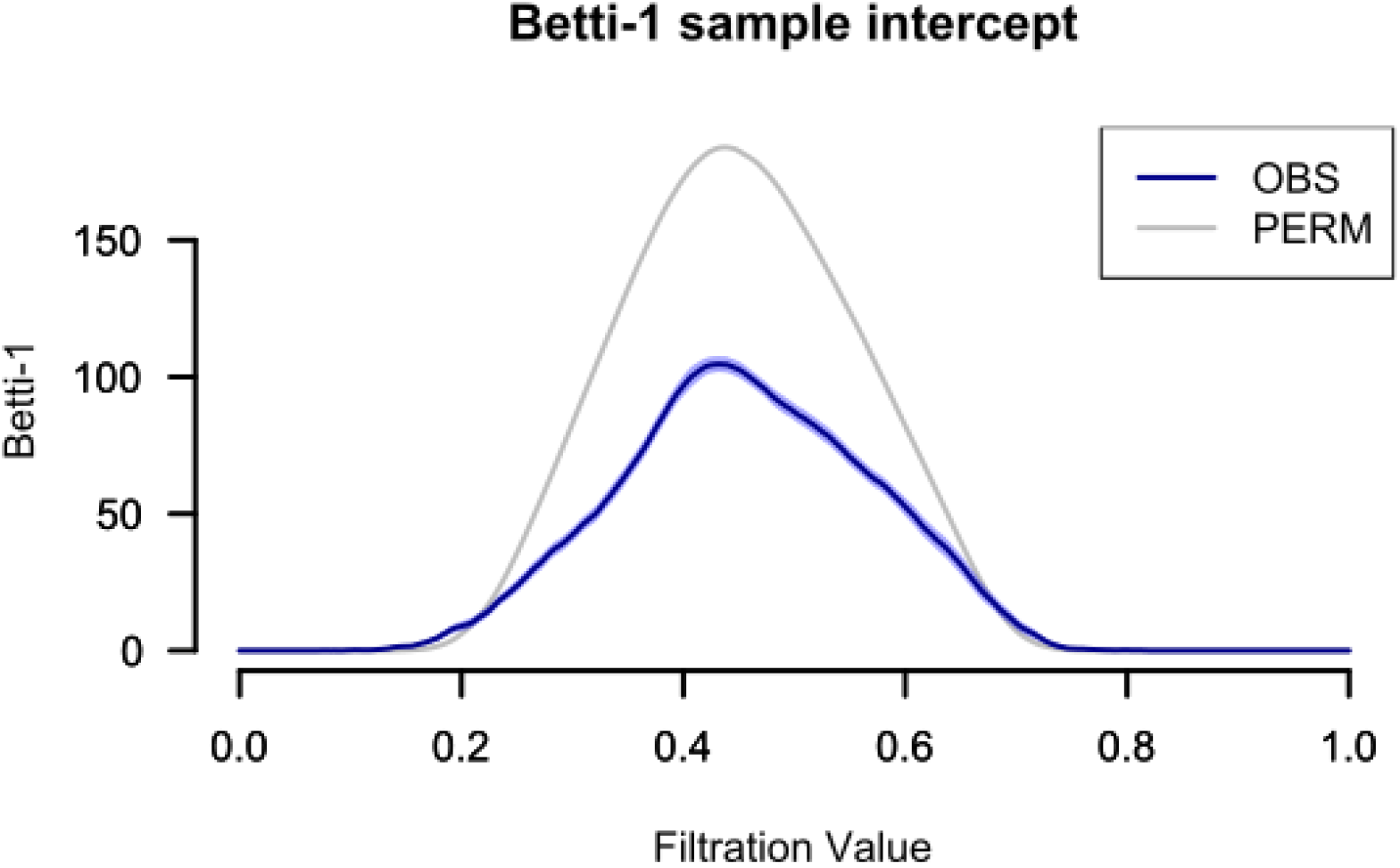
Sample intercept B_1_ curve (with 95% confidence interval) in blue. Average of 1000 bootstrapped connectomes with random edge-rewiring B_1_ curve (with 95% confidence interval) in gray.

The distance between nodes was set as one minus their corresponding Pearson’s correlation (i.e., their functional connectivity edge), following Lee et al. (2012): d(x_i_, x_j_) = 1 - r(x_i_, x_j_). Being *r* the Pearson’s correlation between nodes x_i_ and x_j_. B_0_ and B_1_ curves were computed using the TDA R-package (Fasy et al., 2014), and were summarized by means of the area under the curve (AUC). The AUC accounts for the overall process of the Betti numbers along all possible values of □. Low scores of B_0_-AUC can be interpreted as a fast transition to the single component, while higher scores imply a more segregated configuration of the brain network. Meanwhile, low scores of B_1_-AUC mean that distributed connected components rapidly bind to one another, and higher scores imply an increase in the number of holes within the network (i.e., a more distributed connectivity structure).

Furthermore, to discard that the observed results can be obtained by chance, a null distribution of Betti curves was generated by bootstrapping 1000 connectomes extracted from the original sample whose edges were randomly rewired (Giusti et al., 2015; Gracia-Tabuenca et al., 2020).

### 2.7 Developmental trajectories

Developmental effects were tested using linear mixed-effects (LME) and nonlinear generalized additive mixed models (GAMM). Six models were applied: two LME for age and age-sex interaction, two GAMM fitting smooth splines for age and age-by-sex. However, given that PDS is an ordinal and not a continous scale, nonparametric locally estimated scatterplot smoothing (LOESS) terms for PDS and PDS-sex interaction were included in the two remaining GAMMs. Every model included random-effects for the intercept plus average head motion (FD-RMS) and coil as confounds. Random-effects were estimated via maximum likelihood. Models were implemented using R libraries: LME via *lme4* (Bates et al., 2007; RRID:SCR_015654), GAMM with splines via *gamm4* (Wood et al., 2017), GAMM with LOESS via *gamlss* (Stasinopoulos and Rigby, 2007). Model selection was set by the lowest Akaike Information Criterion (AIC; Akaike, 1974). The AIC evals a model by the tradeoff between its complexity and its goodness of fit. That is, the subtraction between the number of parameters (*k*) and the log-likelihood function (*lnL*) by a factor of two (i.e., AIC = 2*k*-2*lnL*).

In addition, developmental effects within the model were tested using a “drop-term” Likelihood Ratio Test (LRT) approach. The LRT contrasts the full model in relation to a null model without the term of interest. Also, LRT was applied for the thirteen functional networks of the Power et al. (2011) segmentation, where their corresponding significance was corrected for multiple testing using a False Discovery Rate (FDR) q < 0.05 (Benjamini and Hochberg, 1995).

### 2.8 Code accessibility

All preprocessed data and the code described in this study are freely available online at ###. Also, the code is available as Extended Data 1. Present results were computed with an Intel Core i7-4790 CPU @ 3.60□GHz□×□8 with Ubuntu 18.04.3 LTS 64-bit.

## 3. Results

The sample intercept B_0_ curve, that is, the representative curve for the whole sample, showed an inverse sigmoid pattern with a slower transition to the single component compared to the permuted data (Figure 2). On the other hand, the B_1_ curve shows a bell-shape with a maximum of 104.69 “holes” at 0.432 filtration value, while the permuted data shows a maximum of 183.92 at □ = 0.438 (Figure 3).

Regarding model selection for the developmental effects, the GAMM for the PDS showed the lowest AIC for B_0_-AUC (834.83) and B_1_-AUC (615.71) (Table 1).

**Table 1.**
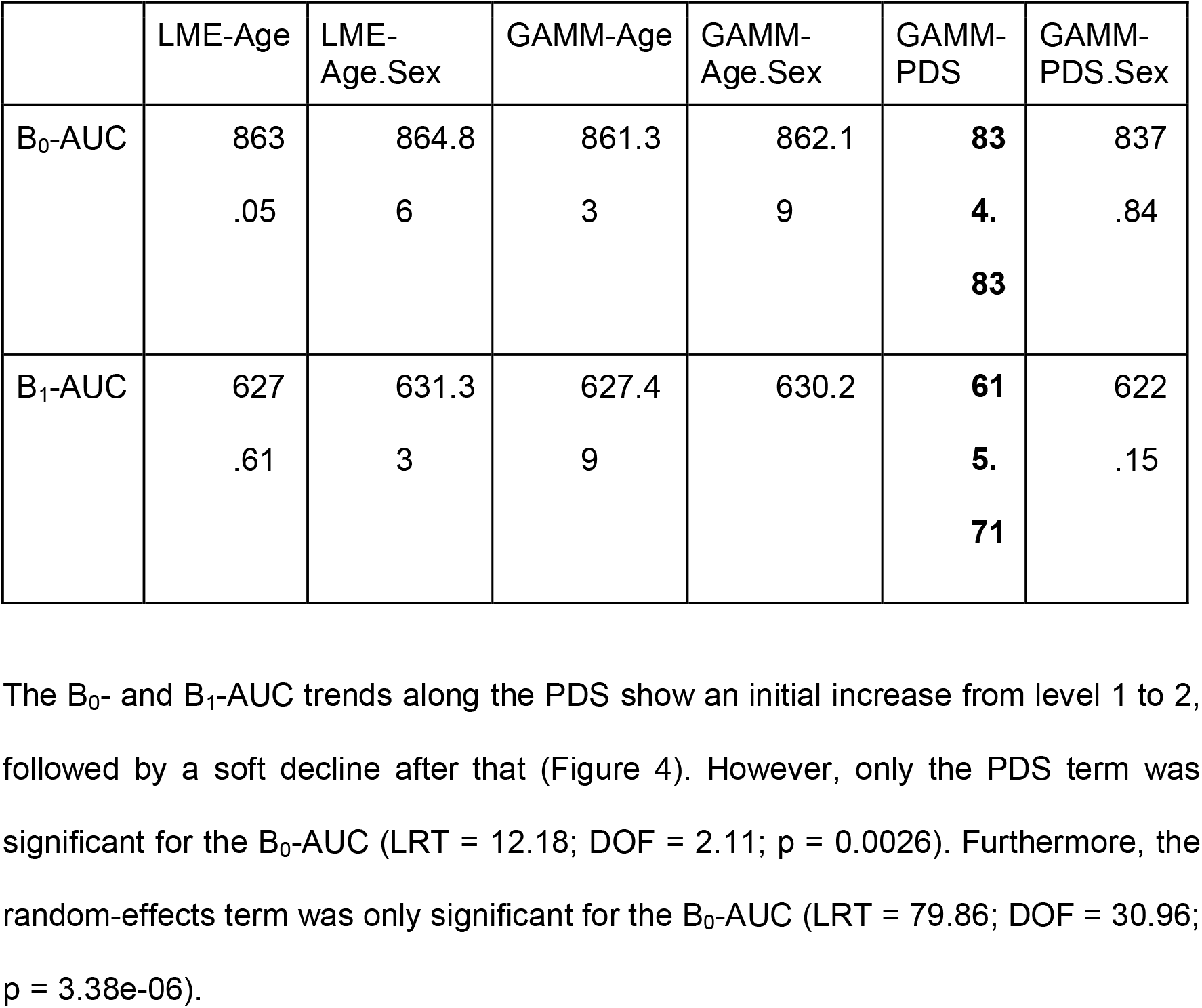
Akaike Information Criterion (AIC) for Betti-0 (B_0_) and Betti-1 (B_1_) areas under the curve (AUC) at every developmental model: linear mixed-effects models for age (LME-Age) and age-sex interaction (LME-Age.Sex), generalized additive mixed models with smooth splines for age (GAMM-Age) and age-by-sex (GAMM-Age.Sex), and with LOESS terms for PDS (GAMM-PDS) and PDS-sex interaction (GAMM-PDS.Sex). The smallest values in bold.

|  | LME-Age | LME-Age.Sex | GAMM-Age | GAMM-Age.Sex | GAMM-PDS | GAMM-PDS.Sex |
| --- | --- | --- | --- | --- | --- | --- |
| B<sub>0</sub>-AUC | 863<br>.05 | 864.8<br>6 | 861.3<br>3 | 862.1<br>9 | **83**<br>**4.**<br>**83** | 837<br>.84 |
| B<sub>1</sub>-AUC | 627<br>.61 | 631.3<br>3 | 627.4<br>9 | 630.2 | **61**<br>**5.**<br>**71** | 622<br>.15 |

Concerning the developmental effects at the functional network level, PDS term showed strong effects in the Fronto-Parietal (FPN) and moderate effects in the Auditory (AUD), sensorimotor-hand (SMH), and subcortical (SUB) networks for the B_0_-AUC (Figure 5). Only the FPN had a significant effect after FDR correction (LRT = 26.18; DOF = 7.21; p = 5.52e-04), which shows a nonlinear trend similar to that for the whole brain network (Figure 5). No effects (even uncorrected) were found for the B_1_-AUC.

**Figure 4.**
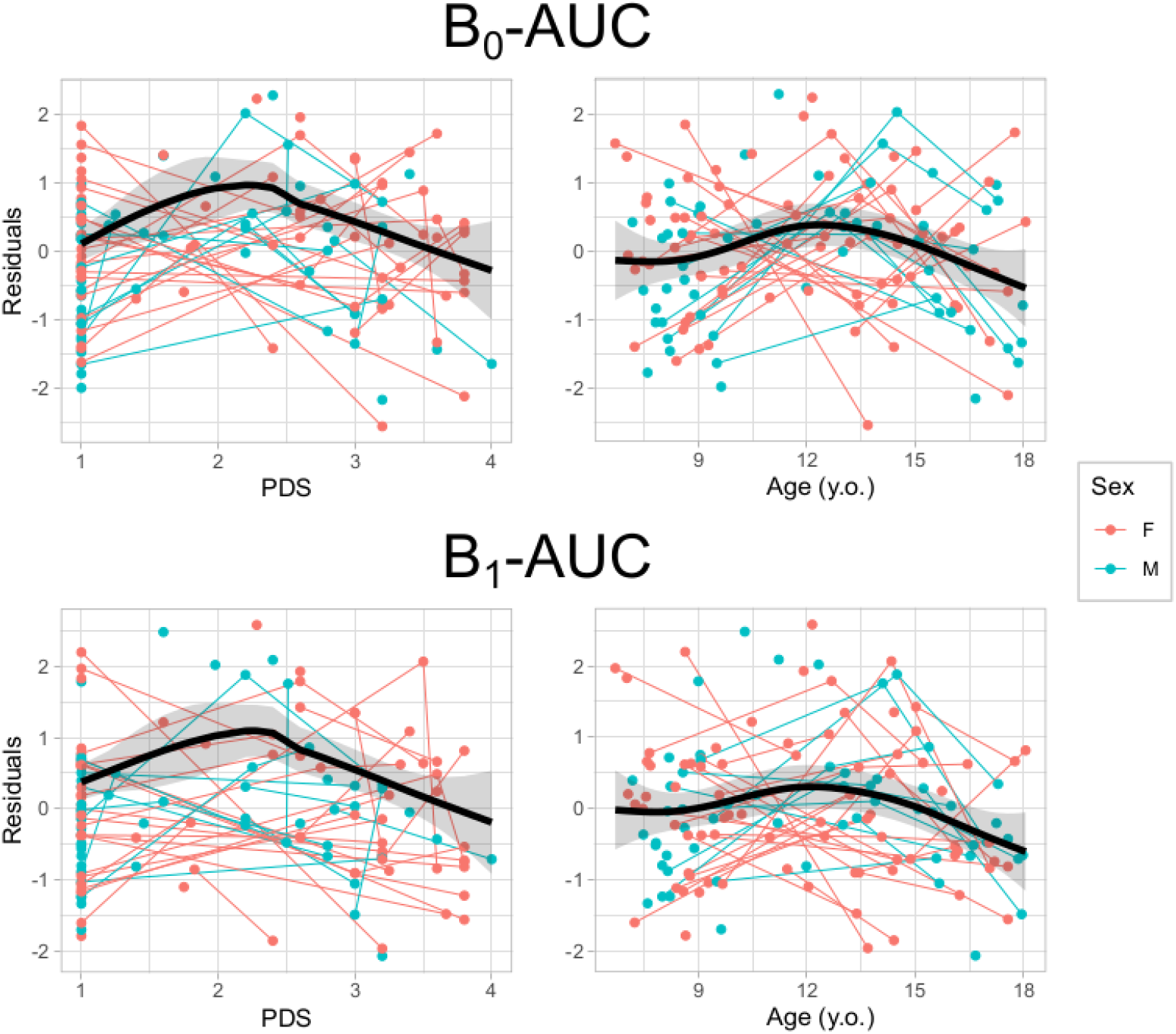
Scatter-plots of the GAMM PDS-LOESS (left) and age-spline (right) models for the B_0_ (top) and B_1_ (bottom) area under de curve (AUC) residuals (after regressing out in-scanner motion and head-coil) in relation to the pubertal scale (PDS) or age. Thin lines represent individual trajectories; thick black lines represent the sample curve (with 95% confidence-interval shadow).

**Figure 5.**
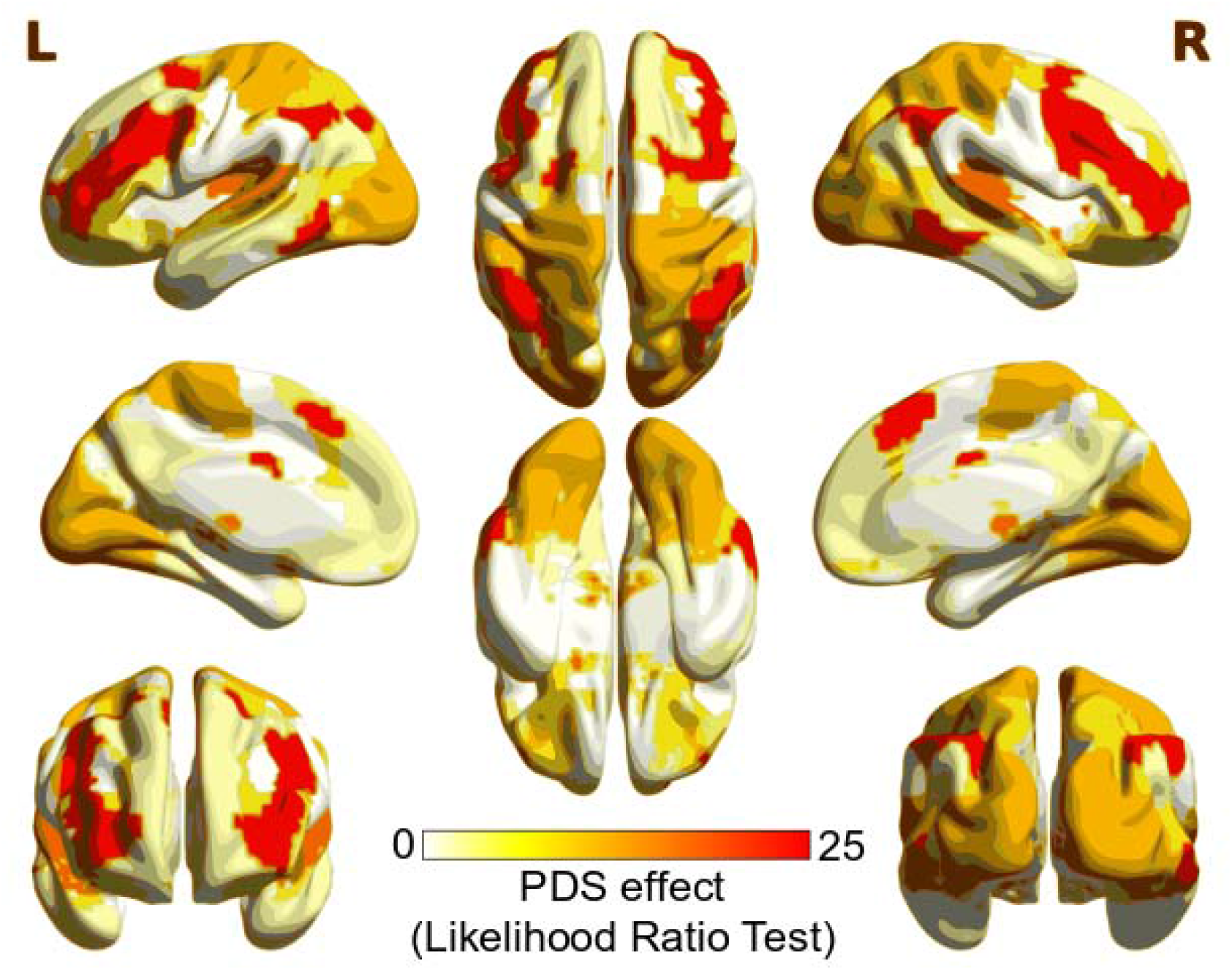
Brain maps of the Likelihood Ratio Test of the GAMM LOESS of the Pubertal Developmental Scale (PDS) term for B_0_-AUC at the functional networks level. Mapping was based on ROI corresponding to the consensus area according to Power et al. (2011), using BrainNet Viewer (Xia et al., 2013; RRID:SCR_009446).

**Figure 6.**
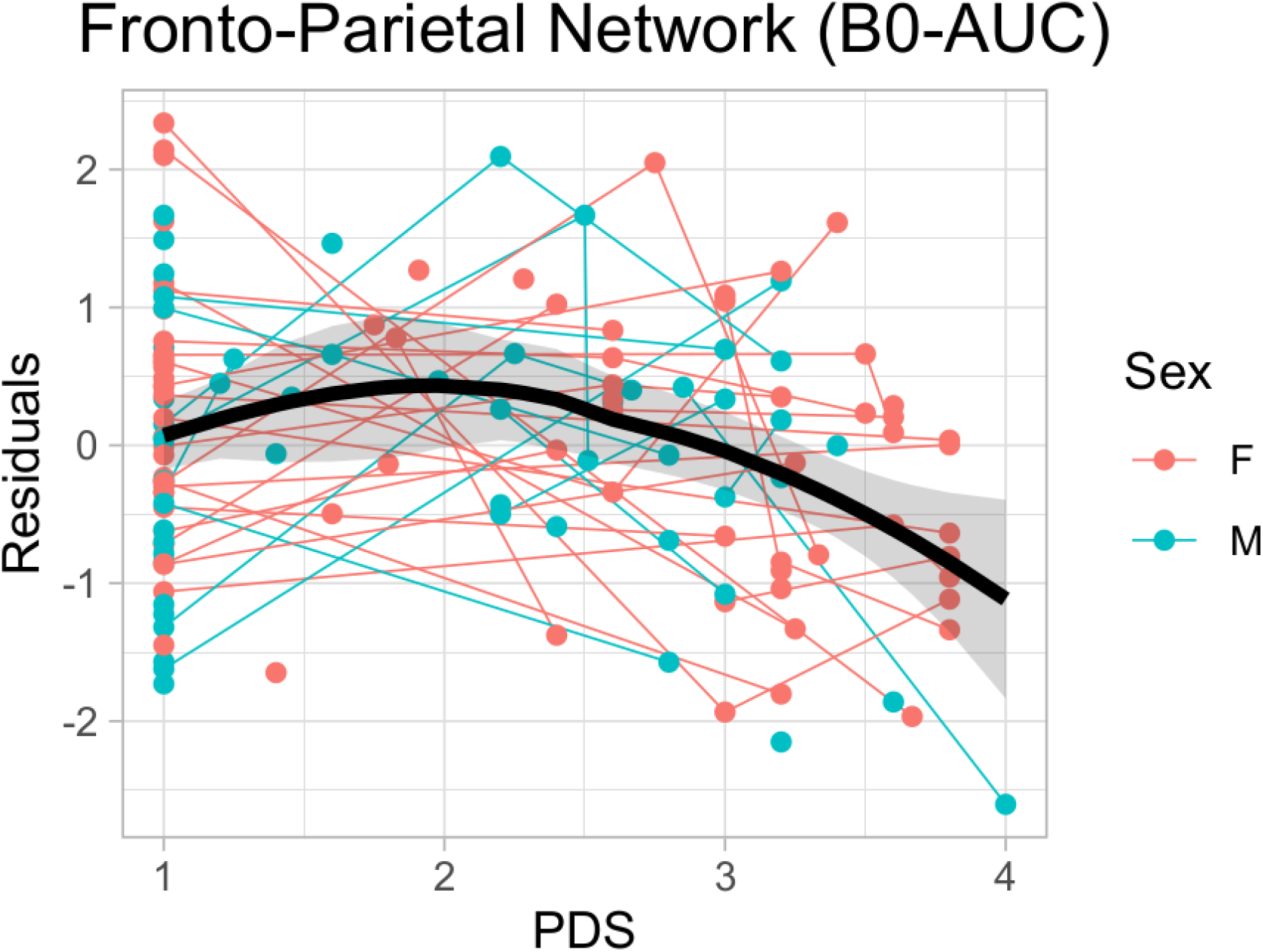
Scatter-plot of the GAMM LOESS-PDS model for the B_0_-AUC residuals (after regressing out in-scanner motion and head-coil) of the Fronto-Parietal Network (FPN) in relation to the pubertal scale (PDS). Thin lines represent individual trajectories; thick black lines represent the sample LOESS curve (with 95% confidence-interval shadow).

## 4. Discussion

In this study, we have applied Topological Data Analysis (TDA) on the functional connectomes of a longitudinal sample of typically developing children and adolescents. TDA features show a segregated connectivity structure compared to random networks, but even segregated, the brain connectomes exhibit a more dense connectivity pattern within neighbours connections. Furthermore, this topology develops in a nonlinear fashion through adolescence, better described by the pubertal status than by chronological age. This nonlinear effect exhibits a faster connectivity of the whole-network and within the Fronto-Parietal Network just after the onset of the pubertal signs.

Regarding the average Betti curves, the sample intercept for the B_0_ curve showed an inverse sigmoid pattern that replicates previous findings in functional connectivity fMRI (Liang and Wang, 2017; Gracia-Tabuenca et al., 2020; Li et al., 2021) and PET (Lee et al., 2012) studies. Furthermore, the average B_0_ curve of the randomized data reached the single component faster than the observed data, evincing a less segregated network. This random pattern was also replicated in another pediatric sample showing the same faster transition to a single component (Gracia-Tabuenca et al., 2020). Concerning the B_1_, both the sample intercept and randomized curves exhibit a bell-shaped curve with an approximate similar filtration value at their maxima, but lower area for the observed data. This evinces a connectivity structure of lower number of holes, or more densely connected at the local level in the real data compared to the random networks. Thus, B_0_ tells how fast the whole-network goes from isolated to all-connected nodes, while B_1_ reflects how densely connected are those elements already connected. It is noticeable that on average B_0_ tends to reach the single component at filtration values of □=0.5, but at that point B_1_ curves display their peak number of “holes”. This implies that these TDA features not only reflect different levels of connectivity structure, but also, they occur at different connectivity strengths.

About the developmental effects of the TDA features, several models were tested to address the area under the B_0_ and B_1_ curves. Nonlinear additive models surpass the goodness of fit of the linear ones, even controlling for the extra number of parameters. Specifically, when considering the pubertal status (assessed by the PDS) without its sex interaction is the model that better fits the AUC for both B_0_ and B_1_. Hence, the development of the functional connectome topology better adjusts the pubertal status than chronological age. This is a relevant finding considering that the pubertal status takes into account non-continuous changes as well as more subtle sex effects than a nonlinear age-sex interaction. No previous studies have addressed the adolescent connectome via TDA, but recent studies have shown a better adjustment with the pubertal status for the developmental trends of the functional connectome (based on graph theory; Gracia-Tabuenca et al., 2021) or the frontostriatal functional connectivity (van Duijvenvoorde et al., 2019).

Concerning whole-brain inferences, the AUC for both B_0_ and B_1_, showed an initial increase from PDS level 1 to 2, but decreased afterwards. In contrast, when focusing on the chronological age, the turning point is approximately at 12 years old, but showing smoother trends compared to PDS. This means a faster transition to the single component for the B_0_, while a lower rate of geometric holes for the B_1_ at the end of the adolescence when considering PDS or age. But only B_0_-AUC effects were significant, which evinces that those changes were more prominent at lower filtration values, i.e., edges with higher functional connectivity (or higher integration). Although TDA was not applied in developmental connectomes yet, previous work on brain functional organization during this period has shown increases along age in the functional segregation (Fair et al., 2009; Satterhwaite et al., 2013a; Gu et al., 2015) and integration (Marek et al., 2015). In addition, when considering the PDS, it has been shown that functional centrality, segregation, efficiency, and integration increases at the end of adolescence (Gracia-Tabuenca et al., 2021). All these studies demonstrate the change in configuration of the brain functional organization during adolescence. Lastly, when testing the random-intercepts effect within the developmental models, in the additive model with PDS for the B_0_-AUC its random-effects showed a strong effect, which shows the relevance of intra-individual trajectories even though of the loss of degrees of freedom.

Likewise, the B_0_-AUC along PDS effect was stronger in the Fronto-Parietal Network (FPN), showing a similar nonlinear trend as that for the whole brain network. This demonstrates a faster integration of the FPN nodes at the end of the adolescence. The FPN is a key module of the connectome that is involved in the response to high-demanding tasks (Zanto and Gazzaley, 2013) and it is a fundamental system for the consolidation of executive behavior in the adolescent period (Baum et al., 2017; Chai et al., 2017). Other works on functional connectomes have reported an increase of the FPN connectivity along with other attention related systems at the late adolescence (Kwang et al., 2013; Marek et al., 2015; Gracia-Tabuenca et al., 2021). Previous studies in animal models have shown brain plasticity associated with puberty-related hormonal changes (Sisk & Foster, 2004). Neuroimaging studies controlling for the age effects, have also revealed structural changes associated with puberty stage in humans, mainly showing decreased gray matter density but increased white matter density in later stages (Peper et al., 2009; Perrin et al., 2009; Bramen et al., 2011; Herting et al., 2012). This work contributes to the emerging evidence that puberty onset greatly influences the development of the brain functional connectivity (van Duijvenvoorde et al., 2019; Gracia-Tabuenca et al., 2021).

Some limitations of this work should be taken into account. We used relatively short scans which may affect the quality of the data, nonetheless it was considered sufficient at the time of the first acquisition (Van Dijk et al., 2010). Furthermore, we applied a strident quality control of the data discarding those datasets with less than 80% good quality data in terms of motion artifact. Nonetheless, the Topological Data Analysis (TDA) complements other network modelling strategies by extracting high-dimensional features across the whole range of connectivity values, instead of exploring a fixed set of connections.

## 5. Conclusion

The present study focused on the characterization of functional connectomes as topological spaces in a longitudinal sample of typically developing children and adolescents. Observed Topological Data Analysis (TDA) features showed higher segregation and denser local connections compared to random networks. However, during adolescence this effect changes with a nonlinear trend that increases the integration of the whole-brain and the Fronto-Parietal Network, particularly after the onset of the pubertal signs. These results provide evidence of the nonlinear, puberty-dependent developmental trajectories of the topology of the brain network. With the advantage that these properties arise exploring the whole range of connectivity strengths instead of focusing on a small set of them. Being adolescence a critical period for the appearance of the first signs of mental health disorders, we expect these trajectories may be of interest for studying both normal and altered development.

